# An assembly-free method of phylogeny reconstruction using short-read sequences from pooled samples without barcodes

**DOI:** 10.1101/2021.04.09.439138

**Authors:** Thomas K. F. Wong, Teng Li, Louis Ranjard, Steven Wu, Jeet Sukumaran, Allen G. Rodrigo

## Abstract

A current strategy for obtaining haplotype information from several individuals involves short-read sequencing of pooled amplicons, where fragments from each individual is identified by a unique DNA barcode. In this paper, we report a new method to recover the phylogeny of haplotypes from short-read sequences obtained using pooled amplicons from a mixture of individuals, without barcoding. The method, AFPhyloMix, accepts an alignment of the mixture of reads against a reference sequence, obtains the single-nucleotide-polymorphisms (SNP) patterns along the alignment, and constructs the phylogenetic tree according to the SNP patterns. AFPhyloMix adopts a Bayesian model of inference to estimates the phylogeny of the haplotypes and their relative frequencies, given that the number of haplotypes is known. In our simulations, AFPhyloMix achieved at least 80% accuracy at recovering the phylogenies and frequencies of the constituent haplotypes, for mixtures with up to 15 haplotypes. AFPhyloMix also worked well on a real data set of kangaroo mitochondrial DNA sequences.

## Introduction

Molecular phylogenetic reconstruction is the mainstay of modern evolutionary biology [1,2]. To use a particularly relevant and recent example, tracing the spread of the COVID-19 pandemic, and understanding the emergence of new variants, has required the use of reliably constructed phylogenies of SARS-CoV-2 genomes [3]. DNA sequencing is used to produce the data from which such valuable phylogenies can be inferred. However, because modern sequencing technologies can produce several gigabases of nucleotide sequences in a single day, one of the challenges for the molecular phylogeneticist is to deal with this quantity of data in a timely manner while still reconstructing accurate phylogenies. To this end, phylogeneticists have developed rapid alignment and tree reconstruction algorithms [4, 5], using pre-processed and curated sequences. Pre-processing and sequence curation can be laborious, but are necessary tasks because a great deal of sequence data are generated using next generation short-read sequencing technologies. Sequences generated in this way are often barcoded using unique DNA identifier tags, and then collectively pooled and sequenced in a single run. The unique barcode allows sequences belonging to different samples to be separated computationally, before additional error-correction and subsequent down-stream analyses are performed.

Quite apart from the costs incurred by data pre-processing and curation, the preparation of barcoded sequence libraries is itself costly. More importantly, there are some samples where barcoding is impractical. For instances, rapidly evolving viruses (e.g., Human Immunodeficiency Virus (HIV) and Hepatitis C Virus (HCV)) typically exist as a collection of genetically diverse genomes within an infected individual. To sequence one or more target genes from a collection of these viruses using barcoding, one would need to isolate individual viral genomes before library preparation. This can be done, but again, is time-consuming and laborious.

In this paper, we describe a novel approach, AFPhyloMix (Assembly-Free Phylogenetics for Mixtures) to reconstruct the phylogeny of non-barcoded amplicons in a mixture that has been sequenced using short-read sequencing. More precisely, the input sample consists of mixtures of anonymous (i.e., non-barcoded) amplicons of a targeted locus, obtained from multiple individuals, each amplicon longer than the read-length of sequenced fragments. We assume that all short-reads can be aligned to the same reference sequence. We have developed our method to work on samples drawn from a population of closely related individuals (i.e., from individuals within a species). In any mixture of individuals drawn from such populations, some amplicons may be identical to others. We refer to a group of identical amplicons as a haplotype [6]. The mixture, therefore, contains a collection of haplotypes, each haplotype being represented by a relative frequency between 0 and 1 (non-inclusive). AFPhyloMix estimates the phylogeny of the haplotypes and their relative frequencies. To validate our approach, we evaluate the efficiency of the method on simulated and real data, and we discuss the conditions under which the method performs well, and its limitations.

## Methods

### Overview

The algorithm, AFPhyloMix, proceeds as follows. Given a mixture of *n* haplotypes, with relative frequencies (*f*_1_, *f*_2_,…,*f_n_*), short-read sequencing generates *k* sequences that can be aligned to a reference sequence. AFPhyloMix then identifies the potential sites with single-nucleotide-polymorphisms (SNPs) from this alignment of reads. Under an infinite-sites model of evolution [7], where each mutation occurs at a new site and any given SNP can have a maximum of two nucleotides, we distinguish between the frequency of a given nucleotide at a given SNP, and the number of SNPs with the same frequency distribution of nucleotides. We refer to these two types of frequencies as the *SNP ratio* and the *SNP frequency,* respectively. For example, assume that in an alignment with three SNPs, site *i* has nucleotides *A* and *G* with frequencies 0.75, 0.25, respectively; site *j* has nucleotides *C* and *T*, with frequencies 0.6 and 0.4, respectively; and site k has nucleotides G and T with frequencies 0.75 and 0.25, respectively. We will adopt the convention of using the smaller nucleotide frequency when identifying the value of a SNP ratio. Therefore, the SNP ratio for site i is 0.25. Sites i and k have the same frequency distribution of nucleotides, even though they may have different constituent nucleotides. In this case, the SNP frequency for the nucleotide distribution instantiated in sites *i* and *k* is 0.67 or 2/3. (We note that the SNP ratios and frequencies are related to the Site Frequency Spectrum [8]; however, because coverage of short-reads vary across the alignment, nucleotide frequencies at each SNP vary as a continuous rational variable rather than as an integer).

In AFPhyloMix, a likelihood function computes the probability of observing the distributions of SNP ratios (data, *D*) along the alignment given their expected distributions, which is itself conditional on a specified tree topology, haplotype frequencies, and sequencing error, assuming an infinite-sites model of evolution. A Bayesian approach is used to compute the posterior probability *P*(*F, T, e*|*D*) of a set of parameters: the frequencies of haplotypes (*F*), the tree topology (*T*), and the sequencing error (*e*), given the observed pattern of the data (*D*), as follows.

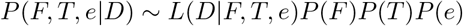

*L*(*D*|*F, T, e*) is the likelihood of the observed pattern of SNPs given the frequencies of haplotypes, tree topology, and the sequencing error. *P*(*F*), *P*(*T*), and *P*(*e*) are the prior probabilities of the frequencies of haplotypes, tree topology, and the sequencing error, respectively. A Bayesian Metropolis-coupled Markov chain Monte Carlo (MCMCMC) inference engine is implemented, to deliver the joint posterior probability distribution of tree topologies and haplotype frequencies. After the Bayesian computation, based on the tree topology with the highest posterior probability, the edge lengths on the tree are computed according to the SNP frequencies.

To illustrate this approach, consider Fig 1 which shows the relationship between observed and expected SNP ratios and frequencies, along a specified tree. Given a 5-tip tree with tip frequencies (i.e. the frequencies of the corresponding haplotypes represented by the tips) shown in Fig 1A, a mutation *x* ∈ {*A, C, G, T*} that occurs on the edge XA over evolutionary time would lead to a different nucleotide on a SNP site in haplotype A relative to other haplotypes. The expected SNP ratio of any mutation along the edge XA would be 0.075, which is the frequency of tip A. The number of SNPs with this mutational pattern — the SNP frequency — would depend on the length of the edge XA. Figure 1B shows the SNP ratio and SNP frequencies and the expected ratio of the occurrences for the mutations on different edges of the tree. For example, the high expected SNP frequency of sites with SNP ratio of 0.485 is due to the mutation on the long edge XZ.

**Fig 1.**
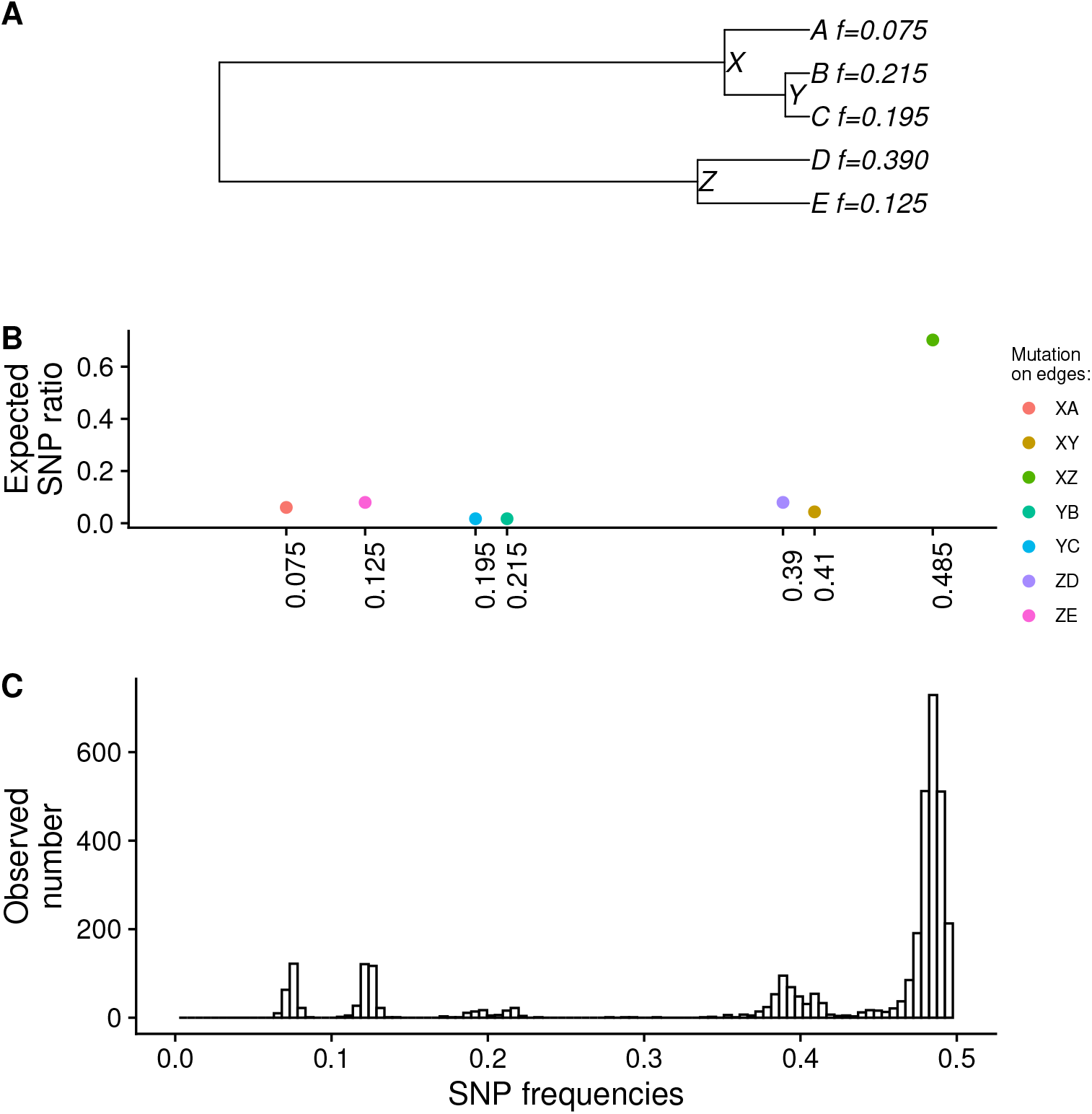
Distribution of SNP frequencies along the genome. (A) An example of 5-tip tree with tip frequencies (i.e. haplotype frequencies). (B) The expected SNP frequencies and the expected SNP ratio of the occurrences for the mutations on different edges of the tree. (C) The observed distribution of SNP frequencies from the short read sequences generated from five simulated genomic sequences with various frequencies based on the tree and the tip frequencies in (A).

### Consideration of the connection between two SNP sites

In Figs 1B and 1C, every SNP site is treated independently. The fact that reads cover multiple sites means that the observed frequencies for multiple sites are correlated. We found that modelling this correlation improved the accuracy of the estimation on the tree topology and the tip frequencies. Where there is no sequencing error, as illustrated in Fig 2A, two SNP sites likely generate three different combinations (*patterns of nucleotides*) on the nucleotide sequences if the mutations of two SNP sites occur on different edges of the tree, while there are only two *patterns of nucleotides* if their mutations happen on the same edge of the tree. For example, two SNP sites with one mutation on the edge ZE and another on the edge XY, as shown in Fig 2A, lead to three different *patterns of nucleotides* on these two SNP sites with expected frequencies 0.125, 0.41, and 0.465. Different locations of the mutations on the tree can result in different sets of expected frequencies (Fig 2B). Similar to the compatibility problem of two sets of binary characters [9], since the infinite site model only allows one mutation along the tree for every SNP site, two SNP sites can create either two or three, but not four *patterns of nucleotides.* Moreover, how often these *patterns of nucleotides* happen depends on the product of the lengths of the edges on which the two SNP sites occur.

**Fig 2.**
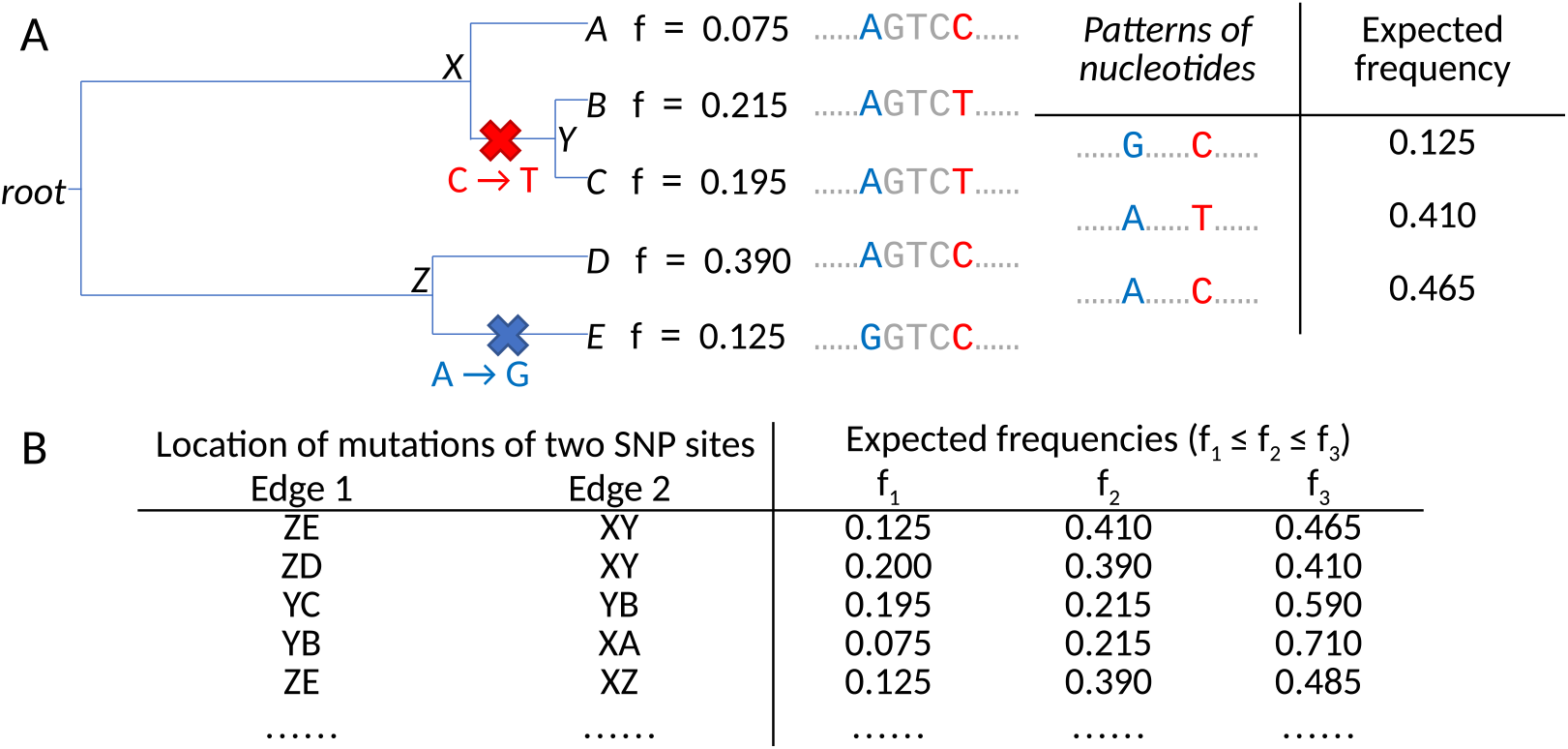
Consideration of connections between two SNP sites. (A) Under the infinite site model, by allowing one mutation along the tree for every SNP site, two SNP sites may make three different *patterns of nucleotides.* (B) Different locations of the mutations on the tree can result in different set of expected frequencies.

Considering the possible three *patterns of nucleotides* (with frequencies *f*_1_, *f*_2_, *f*_3_ = 1 – *f*_1_ – *f*_2_ where *f*_1_ ≤ *f*_2_ ≤ *f*_3_) due to the mutations of two SNP sites on different edges of the tree, Fig 3A shows the distribution of all possible pairs of *f*_1_ and *f*_2_ according to the tree in Fig 2A. The size of the circle represents the expected probability of occurrence. For example, the largest circle at (*f*_1_ = 0.125, *f*_2_ = 0.39) refers to the *patterns of nucleotides* created by two SNP sites with mutations on the edges XZ and ZE, and the probability is relatively high because of their long edge lengths. When we examined the short read sequences generated from five simulated genomic sequences with frequencies based on the tree and the tip frequencies in Fig 2A, we checked every pair of SNP sites close enough to be covered by the same short reads, and obtained the pair of observed values of *f*_1_ and *f*_2_ based on the set of short reads covering the pair of SNP sites. Fig 3B displays the distribution of pairs of observed values of *f*_1_ and *f*_2_. The distribution matches the expected distribution in Fig 3A. Moreover, the distributions of the observed frequencies from the two *patterns of nucleotides* generated by the mutations of two SNP sites on the same edge of the tree were also found consistent with the corresponding expected distributions, although they are not shown here. We are applying a Bayesian approach to the problem and AFPhyloMix estimates the posterior probability distribution of tree topologies and tip frequencies given the observed frequencies of the *patterns of nucleotides* created by pairs of SNP sites.

**Fig 3.**
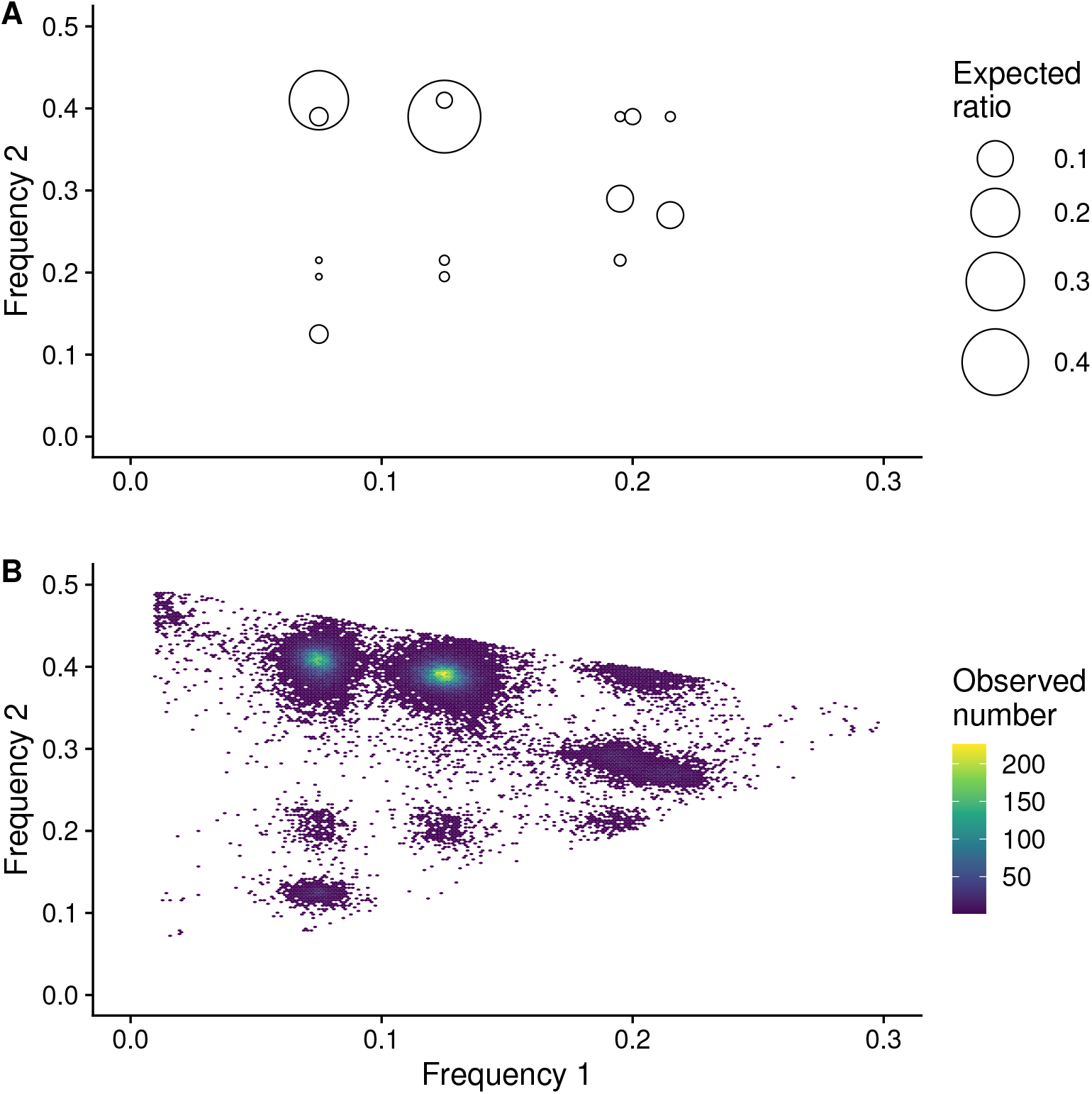
Distribution of the first and the second frequencies of three *patterns of nucleotides* created by two SNP sites. Considering the tree in Fig 2A, two SNP sites having mutations on different pair of edges can lead to three *patterns of nucleotides* with frequencies *f*_1_, *f*_2_, *f*_3_ = 1 – *f*_1_ – *f*_2_, where *f*_1_ ≤ *f*_2_ ≤ *f*_3_. (A) The distribution of all possible pairs of *f*_1_ and *f*_2_. The size of the circle represents the expected chance of occurrence. (B) The distribution of pairs of observed values of *f*_1_ and *f*_2_ obtained from the short read sequences generated from five simulated genomic sequences with frequencies based on the same tree and the same tip frequencies. Every pair of SNP sites close enough to be covered by the same short reads were checked.

### Estimation of tree topology and tip frequencies

Assume that there are n haplotypes. If there is no sequencing error, there should be only two types of nucleotides on each SNP site; for convenience, we will refer to the two allowable states at a given SNP location canonically as ‘0’ and ‘1’. Considering two sites *i* and *j*, let *s*(*ij*) be the nucleotides of the same read covering the sites *i* and *j*. Also let the states of the root of the tree be *r_i_* and *r_j_*, where *r_i_*, *r_j_* ∈ {0,1}, on the site *i* and the site *j*, respectively. Given a *n*-tip rooted tree topology *T*, a set of *n* tip frequencies *F*, and the edges of *T*: {*e*_1_,⋯, *e*_2*n*−2_}, let 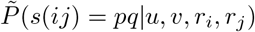, where *p, q* ∈ {0,1} and *u,v* ∈ {*ε,e*_1_,⋯, *e*_2*n*−2_} (the empty string *ε* represents no mutation on the site), be the expected probability of the same sequence having nucleotide *p* on the site *i* and nucleotide *q* on the site *j* given the mutations of the site *i* and *j* are on the edge *u* and the edge *v*, and the states of the root of the tree are *r_i_* and *r_j_*. For example, for the topology and tip frequencies in Fig 2, when *u* = *XY,v* = *ZE, r_i_* = *r_j_* = 0, 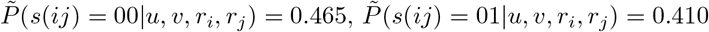, 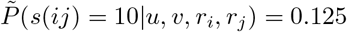 and 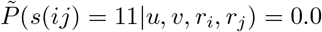.

Ideally, if there is no sequencing error, the number of combinations between the nucleotides of the reads covering the sites *i* and *j* should either be one (if *u* = *v* = *ε*), two (for example, when *u* = *v* ≠ *ε*, or *v* ≠ *u* = *ε*, or *u* ≠ *v* = *ε*), or three. However, in a data set with sequencing error, the number of combinations observed may well be more (up to a maximum of 16). We will compute the expected probabilities taking account of sequencing errors. With the sequencing errors, each SNP site may contain 4 nucleotide types, say 0, 1, 2, and 3. Without loss of generality, we assume 0 and 1 are the major characters, while 2 and 3 are the characters created by the sequencing errors. Given a tree topology *T*, a set of tip frequencies *F*, and a sequencing error rate *e*, define *P*(*s*(*ij*) = *pq*|*u, v, r_i_, r_j_*), where *p,q* ∈ {0,1, 2, 3}, as the expected probability of observing the same read having nucleotide *p* on the site *i* and nucleotide *q* on the site *j*, when the mutations of the sites *i* and *j* are on the edges *u* and *v*, and the states of the root of the tree are *r_i_* and *r_j_*, respectively.

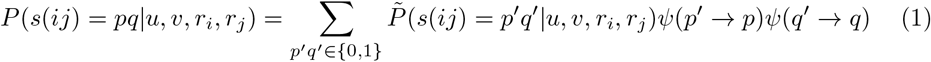

*ψ*(*p*′ → *p*) where *p*′ ∈ {0,1} and *p* ∈ {0,1,2,3} is the proability of observing a nucleotide *p* on the read when the actual nucleotide should be *p*′.

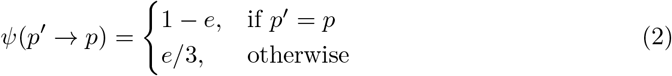

Again, consider two sites *i* and *j*, and let *n_ij_* (*p, q*), where *p,q* ∈ {0,1, 2, 3} be the number of reads observed having nucleotide *p* on site *i* and nucleotide *q* on site *j* in the data set. Let *A_ij_* = {*n_ij_*(*p, q*)|*p, q* ∈ {0,1, 2, 3}} be the observed combinations of characters on the reads covering the site *i* and the site *c_k_*. Given a *n*-tip rooted tree topology *T*, a set of *n* tip frequencies *F*, and a sequencing error rate *e*, define *L*(*A_jj_* |*u, v, r_i_, r_j_, T, F, e*) as the likelihood function of the alignment with sites *i* and *j*, where *i* ≠ *j*, provided that the mutations of the SNP sites *i* and *j* are on the edges *u* and *v*, and the states of the root of the tree on the SNP sites *i* and *j* are *r_i_* and *r_j_*, respectively. We assume that the ratios of the reads having different *patterns of nucleotides* for the sites *i* and *j* follow the multinomial distribution.

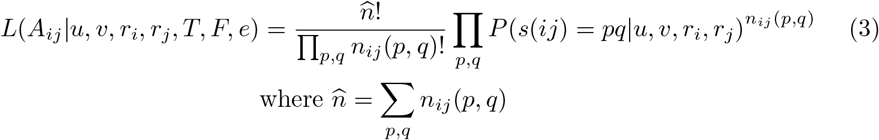

Practically, when performing an analysis on the alignment of the reads, for each site of the alignment, AFPhyloMix assigns the nucleotide supported by the largest number of reads to 0, the second largest to 1, the third largest to 2, and the one with the least supports to 3. There are three reasons to observe two or more nucleotides at a site:

- A site truly has a single mutational event only in its evolutionary history, sequencing errors and other technical artifacts can introduce more than two nucleotides in the alignment of short-reads;
- A site is truly invariable over the evolutionary tree, but sequencing errors/artifacts introduce more than a single observed nucleotide in the alignment at that site; or
- A site truly has experienced multiple mutational events in its events in its evolutionary history (and thus, violates the assumption of an infinite sites model).

We deal with the second and third of these cases below, but if a site truly has only a single mutational event in its history, then nucleotides 0 and 1 should dominate, while nucleotides 2 and 3 will be due to sequencing errors.

Let the *n* SNP sites be {*S*_1_, *S*_2_, *S*_3_, *S*_4_, *S*_5_, *S*_6_,⋯, *S_n_*}. One approach is to consider the patterns observed with pairs of adjacent SNP sites [i.e., if *n* is even, then consider (*S*_1_, *S*_2_)(*S*_3_, *S*_4_) ⋯, (*S*_*n*−1_, *S_n_*))]. This approach allows, at most, only *n*/2 pairs of SNP sites to be considered. On the other hand, if we nominate a reference site, and pair each non-reference site with the reference, we can use ≈ n pairs of SNP sites. We have used this approach, as follows: the whole alignment is partitioned into *m* non-overlapping windows (*W*_1_, *W*_2_, ⋯, *W_k_*, ⋯, *W_m_*) of size *d* (*d* was set to 100 in our implementation). In each window *W_k_* a reference position *c*_k_ ∈ *W_k_* is selected. Let the average of the read coverage along the alignment be *cov_avg_*. The reference site is selected arbitrarily among those sites covered by at least max{50, *r* * *cov_avg_*} reads (where r was set to 0.2). Thus, the selected reference sites will have reasonably high levels of support. For every site i inside the window *W_k_*, its association with the reference position *c_k_* is considered. This approach allows us to consider *n* – *m* pairs of SNP sites. Note that if such reference positions cannot be found (because the coverage of the whole window is not high enough), then a reference covered by the highest number of reads is selected for the window.

The likelihood of the whole alignment (*A*) given the tree topology (*T*), the tip frequencies (*F*) and the sequencing error rate (*e*) is:

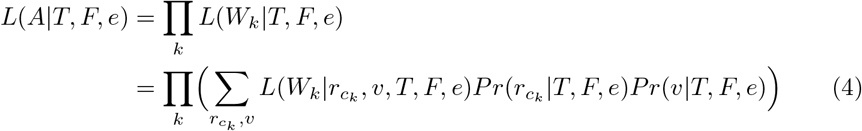

The patterns obtained from the pair of site *i* and the reference *c_k_* depends on the tree topology, the tip frequencies, the sequencing error rate, the root states, and the edges on which the mutations occur for both the site *i* and the reference *c_k_*. Amongst all sites paired with the reference site, pattern frequencies are independent after conditioning on the tree topology, the tip frequencies, the sequencing error rate, the root state and the edge on which the mutation occurs for the reference *c_k_*.

Therefore, the likelihood of the window (*W_k_*) given the tree topology (*T*), the tip frequencies (*F*), the sequencing error (*e*), the root state (*r_c_k__*) and the edge on which the mutation occurs (*v*) for the reference *c_k_* is:

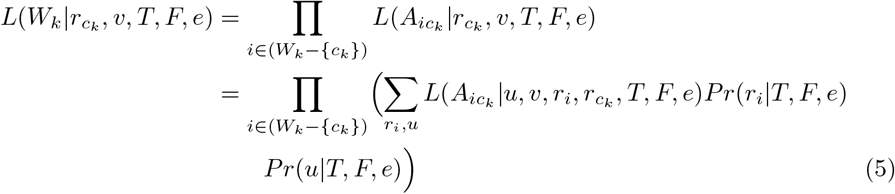

Using this approach, the likelihood essentially integrates across all observed patterns, thus negating the need to identify observed nucleotide ratios that are the consequence of sequencing errors.

During the experiments, the following modified formula was found to have a better accuracy on the estimation of the tree topology and the tip frequencies:

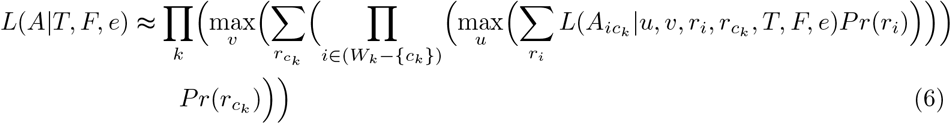

where *P_r_*(*r_i_*) = the observed frequency of the nucleotide *r_i_* in *A*

Define *P*(*T, F, e*|*A*) as the posterior probability of the tree topology *T*, the tip frequencies *F*, and the sequencing error e given the alignment *A*.

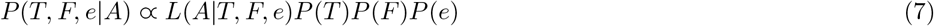

where *P*(*T*), *P*(*F*), *P*(*e*) are the prior probabilities of *T, F, e*, respectively

### Identifying invariable sites

As noted above, a truly invariable site may appear to be a SNP because of sequencing errors. To speed up the computation, AFPhyloMix skips all the sites which are identified as invariable sites. Let the maximum value of the sequencing error rate be *e_max_* (in our application of AFPhyloMix, *e_max_* is set to 0.01). A site is regarded as an invariable site if there exists only one nucleotide such that the percentage of the reads having that nucleotide covering the site is higher than *e_max_*. Let *I* be the set of these sites which are identified as invariable sites; then

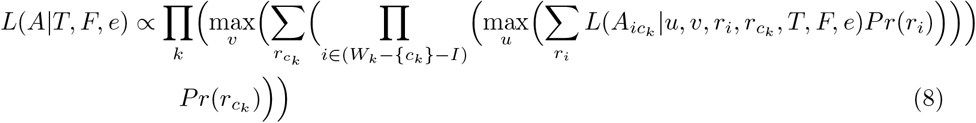

### Markov chain Monte Carlo implementation

AFPhyloMix adopts a Bayesian model of inference to obtain an estimate of the joint posterior probability of phylogenies, haplotype frequencies, and sequencing error, using Markov chain Monte Carlo (MCMC) sampling [10,11]. The MCMC approach has been extensively used in phylogenetic analysis [12,13], but sampling chains may not mix as well as they should. To overcome this, the Metropolis-coupled Markov chain Monte Carlo (MCMCMC) approach was developed by [14]. Although MCMCMC requires multiple parallel sampling chains to be run simultaneously (and thus, has demanding computational overheads), the approach has been demonstrated to improve mixing and convergence to a stationary posterior probability distribution [15]. We implemented MCMCMC in AFPhyloMix, which reports a the tree topology and the tip frequencies which gives the highest value among all the resulting posterior probabilities along the computation.

### Estimation on edge lengths

After AFPhyloMix estimates the topology *T*, the tip frequencies *F*, and the sequencing error *e* for the mixture of short read sequences from *n* haplotypes, AFPhyloMix calculates the edge lengths in *T* (with edges *e*_1_,⋯, *e*_2*n*−2_) by the following method.

Let *length*(*u*) be the length of edge *e_u_*. In AFPhyloMix, *length*(*u*) is approximated as the probability of having mutation on edge *u* along the tree. Note that *length*(*u* = 0) is the probability of no mutation along the tree (i.e. ∑_0≤*u*≤2*n*−2_ *length*(*u*) = 1).

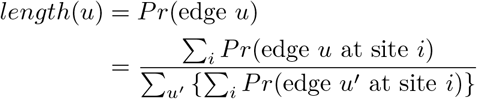

where *Pr*(edge *u* at site *i*) is the probability of a mutation on edge *u* of the tree at site *i*.

As described above, the whole genome is partitioned into non-overlapping windows (*W*_1_, *W*_2_, ⋯, *W_k_*, ⋯) of size *d* (*d* was set to 100), and inside each window *W_k_* a reference site *c_k_* ∈ *W_k_* is selected. For every site *i* (say it is inside the window *W_k_*), we consider the connection between the site *i* and the reference position *c_k_*. For *i* = *c_k_*, let *n_ic_k__*(*p, q*), where *p, q* ∈ {0,1, 2, 3}, be the number of reads observed having nucleotide *p* at site *i* and nucleotide *q* at site *c_k_* on the same read. Define *A_ic_k__* = {*n_ic_k__* (*p, q*)|*p, q* ∈ {0,1, 2, 3}} as the observed combinations of characters on the same reads at the sites *i* and *c_k_*. Similarly, for *i* = *c_k_*, let *n_c_k__* (*q*), where *q* ∈ {0,1, 2, 3} be the number of reads observed having nucleotide *q* at site *c_k_*, and let *A_c_k__* = {*n_c_k__* (*q*)|*q* ∈ {0,1, 2, 3}} be the observed pattern of characters on the reads at the site *c_k_*.

To calculate *Pr*(edge *u* at site *i*), two cases are considered:

- Case 1: *i* ≠ *c_k_* (i.e. the site *i* is not the reference site of the window);
- Case 2: *i* = *c_k_* (i.e. the site *i* is exactly the reference site of the window). For case 1,

*Pr*(edge *u* at site *i*)

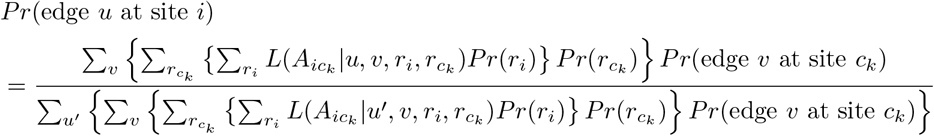

For case 2,

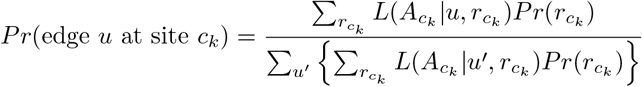

*Pr*(*r_i_*) and *Pr*(*r_c_k__*) are the probabilities of the root states *r_i_* and *r_c_k__*, which are set to the observed frequencies of *r_i_* and *r_c_k__* in *A*, respectively. *L*(*A_ic_k__*|*u, v, r_i_,r_c_k__*) is the likelihood value of the observed combinations of characters on the same reads at the sites *i* and *c_k_* given the mutations on edges *u* and *v* and the root states *r_i_* and *r_c_k__* for the sites *i* and *c_k_*, respectively, while *L*(*A_c_k__*|*u,r_c_k__*) is the likelihood value of the observed pattern of characters on the reads at the site *c_k_* given the mutation on edge *u* and the root state *r_c_k__* for the site *c_k_*. The calculation of *L*(*A_c_k__*|*u,r_c_k__*) can be done by using the similar approach. Whereas it is theoretically possible to simultaneously infer edge lengths and topologies, we have found that this does not deliver accurate results. This is because, in the absence of sequence information at the tips, mutations will naturally favour long branches, thus lowering the probability of seeing short branches. But estimating edge lengths after the topology has been estimated, we overcome this bias. An alternative is to define a suitable prior (e.g., a coalescent prior [16]) that will override the tendency of the likelihood to favour long branches.

### Removal of the sites that violate the infinite sites model

AFPhyloMix uses an infinite site model for modelling the evolution of the genomic sequences between the haplotypes, and thus assumes that every site has no or only one mutation in its evolutionary history. Before AFPhyloMix proceeds to estimate the topology and the tip frequencies, AFPhyloMix examines the read alignments and filters out the sites likely having more than one mutation. The procedure to identify these sites is as follows:

1. If the SNP site has more than two different characters supported by at least *e_max_* (i.e. 1%) of the reads, then the SNP site is ignored.
2. Consider the SNP sites with only two different characters supported by at least 1% of the reads. If there is no back/hidden mutation, two SNP positions will give at most three combinations (as mentioned before). Based on this observation, the following simple method has been developed to identify the sites which are likely to have back/hidden mutations: Consider a SNP site *i*, we check all other SNP sites *j* such that there are sufficient number of reads covering both sites *i* and *j*. If there are at least d sites (i.e. *j*_1_, *j*_2_, ⋯, *j_d_*) which separately have more than three combinations supported by at least 1% of the reads when considering together with site *i*, then the site *i* is regarded having back/hidden mutations and it is discarded. We have tested for different values of *d* on simulated data and found that when *d* = 3 the method performed reasonably well in terms of accuracy.

## Experiment and results

Both simulated and real data were used to evaluate AFPhyloMix.

### Simulated data

Six hundred data sets were simulated and each data set was a mixture of various numbers (*n*) of haplotypes, where *n* ∈ {5, 7, 9,11,13,15} (100 data sets each). In each data set, the n haplotypes were mixed in different random concentrations (with the difference between any two haplotypes ≥ 0.0033). Paired-end reads of length 150bp with total coverage of 15,000x were simulated by ART [17] with the Illumina HiSeq 2500 error model - HS25, from n 50k-long haplotype sequences, which were generated by INDELible [18] using JC model [19] from a *n*-tip tree with 0.03 root-to-tip distance randomly created by Evolver [20] from PAML package [21].

The root sequence reported by INDELible [18] was used as the reference sequence. After using BWA [22] to align the reads against the reference sequence, we ran AFPhyloMix under the default settings on the read alignments to estimate the tree topology and the tip frequencies (i.e. concentrations of the sub-samples) for the mixture. By default, AFPhyloMix runs 8 MCMC processes in parallel: one cold chain and seven hot chains, and each chain runs 650K (for mixture with 5 sub-samples) to 1150K (for 15 sub-samples) iterations, depending on the number of edges in the resulting tree. Fig 4 shows a result from AFPhyloMix on a simulated data set with a mixture of 15 sub-samples. Fig 4A displays the posterior probabilities along the cold chain of MCMCMC process while running the AFPhyloMix. The posterior probabilities increased rapidly during the burn-in period and then appeared to stabilise to an equilibrium distribution. Fig 4B shows the distribution of tip frequencies (i.e. haplotype concentration) along the cold chain of MCMCMC process from AFPhyloMix. The distribution was found to match the expected tip frequencies (marked with red dots) from the real tree used to simulate the data set. Fig 4C is the resulting tree from AFPhyloMix with tip frequencies. This is the tree with the highest posterior probability along the cold chain of MCMCMC process. The tree is topologically congruent with the real tree (Fig 4D) and the tip frequencies are also very similar with the real sub-sample concentrations.

**Fig 4.**
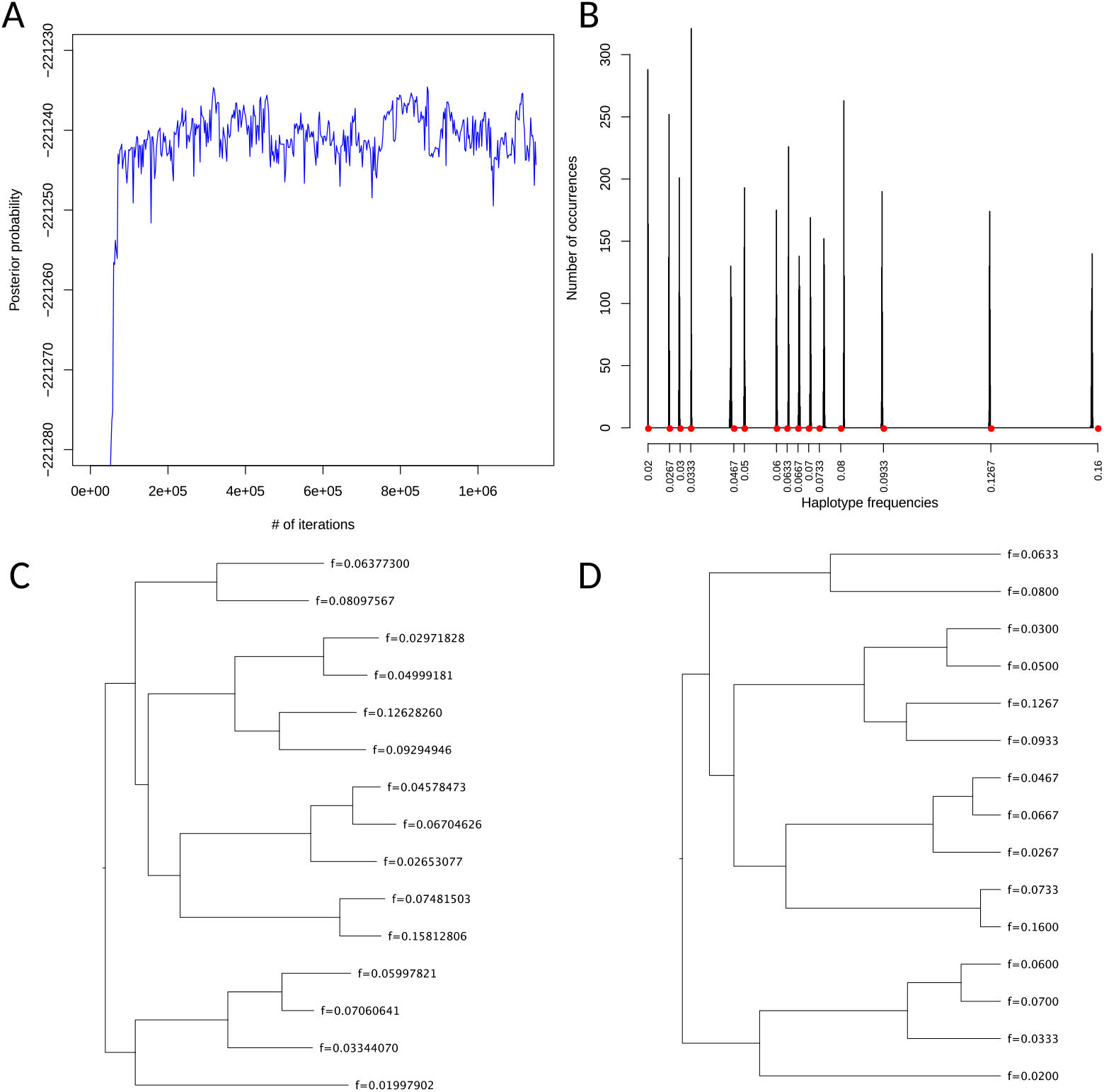
The result of AFPhyloMix on a simulated data set with a mixture consisting of 15 sub-samples. AFPhyloMix was run under the default settings on the read alignments of a simulated mixture of 15 sub-samples in order to estimate the tree topology and the tip frequencies. (A) The posterior probabilities along the cold chain of MCMCMC process while running the AFPhyloMix, which increased rapidly during the burn-in period and then frustrated steadily over a range of values and was reaching a convergence. (B) The distribution of tip frequencies (i.e. sub-sample concentration) along the cold chain of MCMCMC process from AFPhyloMix. The actual sub-sample concentrations are marked by the red dots. (C) The tree with tip frequencies having highest posterior probability along the computation reported by AFPhyloMix. (D) The real tree with the actual sample concentration used for simulating the data set.

Fig 5A shows the summary on the accuracy of AFPhyloMix running on the simulated data sets. Among the data sets with the same number of haplotypes, the figure shows the percentage of data sets with *correct* estimation on both topologies and tip frequencies. The estimated result is regarded as *correct* if the tips on an estimated tree can be paired up with the tips on the actual tree satisfying the following conditions: (1) the difference between the predicted tip frequency and the corresponding actual tip frequency paired with is less than 0.01; and (2) their topologies are the same. From the figure, AFPhyloMix achieved at least 80% accuracy for the mixtures with up to 15 haplotypes. To further examine the derivation between the predicted tip frequencies and the actual sample concentrations, and that between the estimated and the actual edge lengths, for each data set with *correct* estimation, we computed the root-mean-square value of the differences between the estimated and the actual values. Figure 5B and Figure 5C show the summary on the distributions of the root-mean-square of the differences for the tip frequencies and the edge lengths. The values rise gradually as the number of haplotypes increase. For tip frequencies, the root-mean-square values were all below 0.0012, while over 75% of the cases were below 0.0008. For edge lengths, the root-mean-square values were all below 0.005.

**Fig 5.**
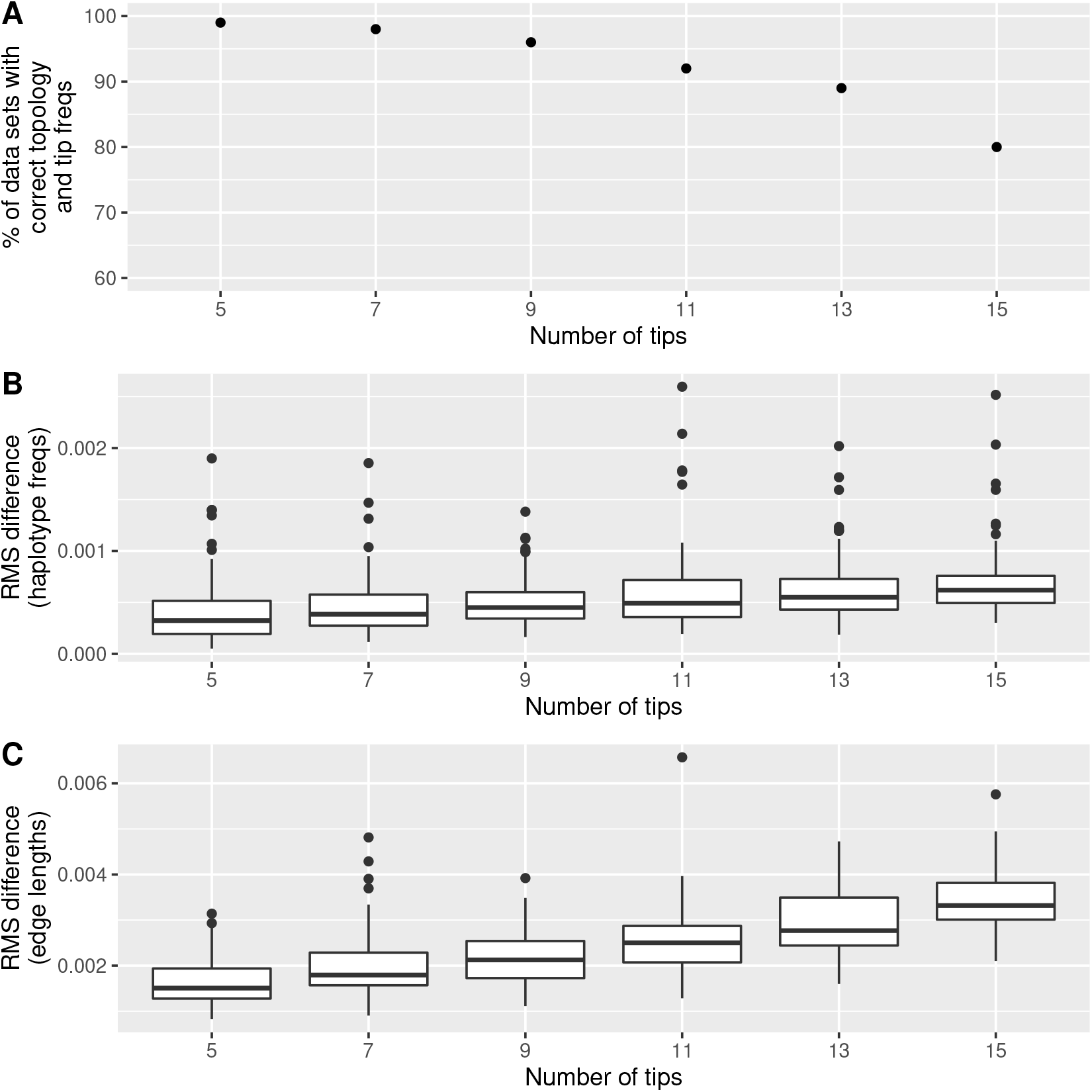
Performance of AFPhyloMix on simulated data. (A) Accuracy of AFPhyloMix - The percentage of data sets (out of 100), for different number of haplotypes, with correct estimated topologies and predicted tip frequencies. (B) Root-mean-square differences between the actual and the predicted tip frequencies. (C) Root-mean-square differences between the actual and the predicated edge lengths.

The pooled 95% highest posterior density of the MCMCMC estimate of error rates on these simulated data sets was between 0.00199 and 0.00204 with an average 0.00202. ART [17] reports the errors for the reads simulated by the error model HS25; in our simulations, we obtained a simulated error rate of 0.00190. AFPhyloMix relies on the alignment of reads against a root sequence to obtain the marginal posterior probability distribution of error rates. We expect, therefore, that the slightly higher value (≈ +0.00012) of the MCMCMC estimate of error rates on these simulated data sets, compared with the simulated error rate reported by ART, is likely due to imperfect alignment between simulated reads and the root sequences.

### Real data

Apart from the simulated data sets, mixtures of reads from kangaroo haplotypes were also used to evaluate the performance of AFPhyloMix.

#### DNA material collection and extraction

In Australia, kangaroo are not farmed, but are culled annually to control population numbers. Culled animals are butchered by certified butchers, and the meat is sold in supermarkets. As the exact provenance of kangaroo meat sold at supermarkets is unknown, meat (produced by Macro Meats) was purchased at local supermarkets in the Australian Capital Territory (Coles Supermarkets Australia Pty Ltd and Woolworths Ltd) on 10 separate occasions over one year (from 29th May 2018 to 26th July 2019), to avoid sampling the same animal. According to statistics from the Department of the Environment and Energy of Australia, they should be of genus *Macropus,* and most likely eastern grey kangaroo, *Macropus giganteus,* as it is the largest population with highest quota in New South Wales, Australia [23].

Approximately 30 mg of meat was excised and homogenized for each individual. Genomic DNA was then extracted using the DNeasy Blood & Tissue Kits (Qiagen) following the manufacturer’s protocol.

#### Amplification and sequencing

Long PCR amplification of complete kangaroo mitochondrial genomes was carried out using the pair of primers (Lt12cons: 5’-GGGATTAGATACCCCACTAT −3’, HtPhe: 5’-CCATCTAAGCATTTTCAGT −3’), which was selected from a previous study [24]. PCR reactions was performed using Takara PrimeSTAR GXL DNA Polymerase under the following conditions: 1 min initial denaturation at 95°C, followed by 30 cycles of 10 s at 98°C, 15 s at 55°C, and 15 min at 68°C. The PCR products were electrophoresed in 1% agarose gel, purified the fragments, and then randomly fragmented to 650 bp by sonication (Covaris S220).

Library preparation and sequencing were performed by GENEWIZ. Amplified fragments of all 10 individuals were sequenced under the same run. In order to obtain a highly reliable phylogenetic tree of these sub-samples for evaluating our method, each individual was barcoded with unique indices before multiplexing and sequencing, so that each short read sequence could be identified to the corresponding sub-sample. The relative concentrations of the sub-samples are listed in the Table 1 (2^*nd*^ column). Sequencing was performed on an Illumina MiSeq machine with paired-end read length of 2 x 300 bp.

**Table 1.** Concentration of kangaroo sub-samples

| Sub-sample ID | Concentration | Concentration without K01 |
| --- | --- | --- |
| K01 | 0.019 | - |
| K02 | 0.029 | 0.030 |
| K03 | 0.045 | 0.046 |
| K04 | 0.047 | 0.048 |
| K05 | 0.094 | 0.096 |
| K06 | 0.102 | 0.104 |
| K07 | 0.125 | 0.127 |
| K08 | 0.130 | 0.132 |
| K09 | 0.188 | 0.192 |
| K10 | 0.221 | 0.225 |
| Total | 1.000 | 1.000 |

The relative concentration between different kangaroo sub-samples.

#### Phylogenetic tree reconstruction

To start with, a reliable phylogenetic tree between the haplotypes was constructed, so that this gold-standard result could be used to evaluate our method. First, all the short read sequences were demultiplexed into sub-samples according to the barcodes appended on the sequences. Then *de-novo* assembly was performed on the short reads for each sub-sample separately by SOAPdenovo-Trans-127mer from SOAPdenovo-Trans package [25] with parameter: kmer-size=91. After the mitochondria DNA sequences of the 10 haplotypes were constructed, a multiple sequences alignment was computed by MAFFT [4] with G-INS-i strategy in Geneious 11.1.5 [26]. The phylogenetic analysis was conducted using IQ-TREE [27] with the evolutionary model HKY+F+I, and the ML phylogenetic tree (shown in Fig 6B) was used as the reference tree to evaluate our method.

**Fig 6.**
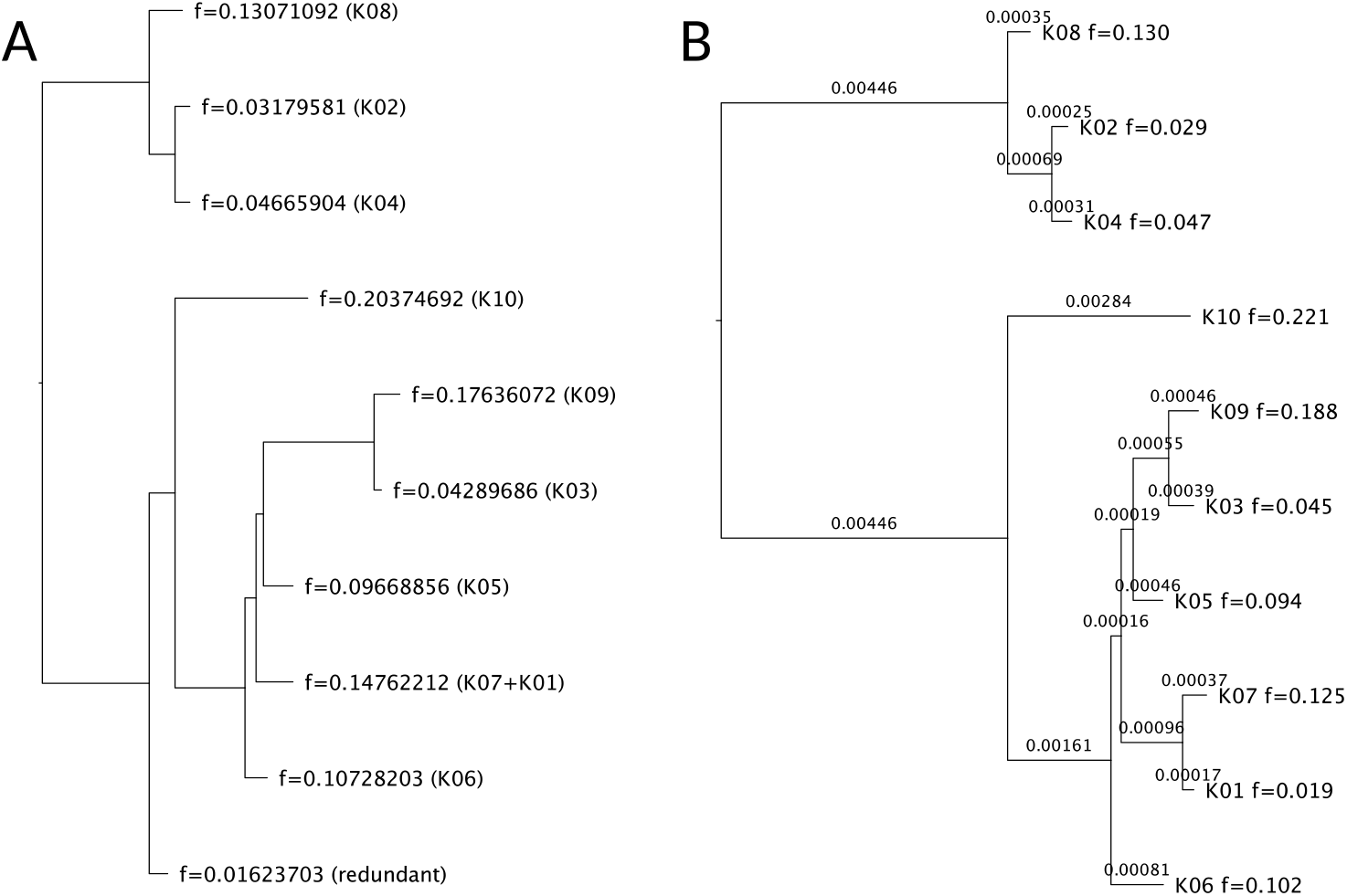
Result on a real data set with mixture of 10 sub-samples. (A) The tree with tip frequencies reported by AFPhyloMix. (B) The tree reported by IQ-Tree [27]. Note that the tip labels inside the brackets in (A) were added manually after comparing with the tree reported by IQ-Tree, indicating the corresponding sample that each tip should be assigned to according to the topology and the tip frequencies.

The short read sequences from 10 haplotypes were then mixed together, and the barcode of each haplotype was removed. Then Trimmomatic [28] was run on the mixture of reads to remove adaptors, leading and trailing low quality bases by using the options: “ILLUMINACLIP:TruSeq3-PE-2.fa:2:30:10 LEADING:3 TRAILING:3 SLIDINGWINDOW:4:15 MINLEN:36”. Then the reads were aligned to the reference sequence of the eastern gray kangaroo mitochondrial genome (GenBank Accession Number: NC_027424) by BWA [22] and the alignments with mapping quality score (MAPQ) lower than 20 were discarded.

AFPhyloMix was used with the short-read alignment to estimate the phylogenetic tree and the relative frequencies of each haplotype. For each read, AFPhyloMix discarded the nucleotides with base quality score lower than 25. Fig 6A shows the reported tree, as well as the tip frequencies, with the highest posterior probability obtained. The tip labels inside the brackets were added manually after comparing with the tree reported by IQ-Tree (in Fig 6B), indicating the corresponding sample that each tip should be assigned according to the topology and the tip frequencies. Overall, the topology and the tip frequencies outputted by AFPhyloMix matched with the tree from IQ-Tree, except that AFPhyloMix combined two haplotypes K01 and K07 into one. From the tree reported by IQ-TREE, the distance between the tips K01 and K07 is 0.00054 (99.946% similarity between these two sequences). The method could not distinguish the two too-similar sequences, and thus regarded them as the same sequence.

To detect whether these two nearly identical sequences affect the performance of AFPhyloMix, we removed the sub-sample of K01 and re-estimated the phylogeny with the same method as described above. The updated relative concentrations of the remaining haplotypes among the mixture is shown in Table 1 (3^*rd*^ column). Both AFPhyloMix and IQ-TREE analyses resulted in the same topology associated with tip frequencies well matched the concentrations of those 9 haplotypes (in Fig 7).

**Fig 7.**
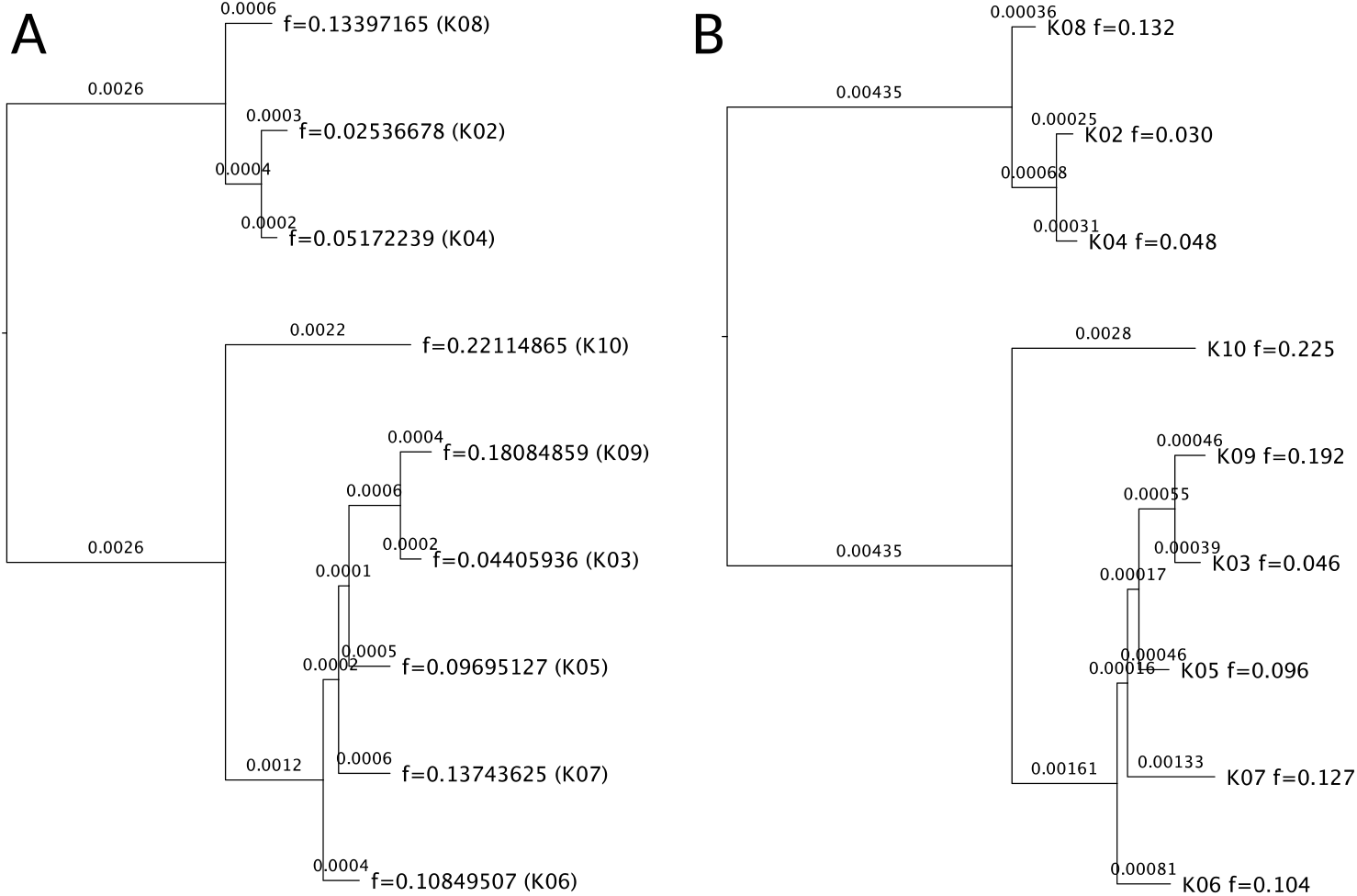
Result on a real data set with mixture of 9 sub-samples. The haplotype K01 was removed from the mixture and the experiment was repeated. (A) The tree with tip frequencies reported by AFPhyloMix. (B) The tree reported by IQ-TREE [27]. Note that the tip labels inside the brackets in (A) were added manually after comparing with the tree reported by IQ-Tree, indicating the corresponding sample that each tip should be assigned to according to the topology and the tip frequencies.

The 95% highest posterior density of the MCMCMC estimate of error rates on the real data sets was between 0.000722 and 0.000735 with an average 0.000729. In order to compute the underlying actual error rate on the real data set, reads which had been processed by Trimmomatic [28], were compared with the corresponding assembled haplotype and the error rate of 0.000713 was obtained, after discarding the read bases with base quality scores lower than 25 (the same criteria AFPhyloMix used to filter out the low-quality read bases). Again, the slightly higher value (≈ +0.000016) of the MCMCMC estimate of error rates on the real data sets compared with the obtained error rate is consistent with what we observed with simulated data, and is likely due to the imperfect alignment between the reads and the reference sequence.

## Discussion

This research demonstrates the feasiblity of reconstructing a phylogenetic tree directly from the short read sequences obtained from a mixture of closely related amplified sequences, without barcoding. The results indicate that our methods work well on the simulated data set for a mixture of reads generated from up to 15 haplotypes and on a real data set of a mixture with 10 haplotypes.

Perhaps unsurprisingly, AFPhyloMix worked better in the simulated data sets than in the real data sets when it came to estimating haplotype concentrations. The root-mean-square difference between the estimated sub-samples concentrations and the expected concentrations in the real data sets was 0.0059 (from Fig 7), which was larger than those in the simulated data sets (i.e. all were below 0.0027 in Fig 5B). Of course, the expected haplotype concentrations may have differed from the true concentrations in the mixture: the physical act of mixing small volumes could have led to differences in the relative concentrations of haplotypes, and this may have contributed to a higher-than-expected root-mean-square.

Another factor affecting the performance of the method is the varying coverage of short read sequences along the genome. Fig 8A shows the actual distribution of the read alignments of 10 haplotypes along the genome. We expected read coverages to vary along the genome randomly, without any association to haplotypes. Surprisingly, we found that, when all the haplotypes were sequenced under the same run (i.e. all haplotypes were pooled into the same library before sequencing), the read coverages of the haplotypes had similar trends: all had relatively high (or low) read coverages at the same regions of the genome. As shown in Fig 8B, the similar trends of the read coverages along the genome between the haplotypes led to a steady distribution of the ratios on the read coverages between the haplotypes along the genome. The consistent read coverage ratios along the genome worked in our method’s favor; on the other hand, as shown in Fig 8B, we noticed that few short regions on the genome had sudden changes in the ratios of the read coverage amongst haplotypes. In those regions, some reads were trimmed after the alignment against the reference sequence or could not be aligned to the reference sequence due to the dissimilarity between the read sequences and the reference sequence. Another preprocessing step was therefore developed in AFPhyloMix, in which read alignments were examined and problematic regions removed if there existed over *r* (*r* was set to 2% of the read coverage) reads being trimmed around the same sites on the genome.

**Fig 8.**
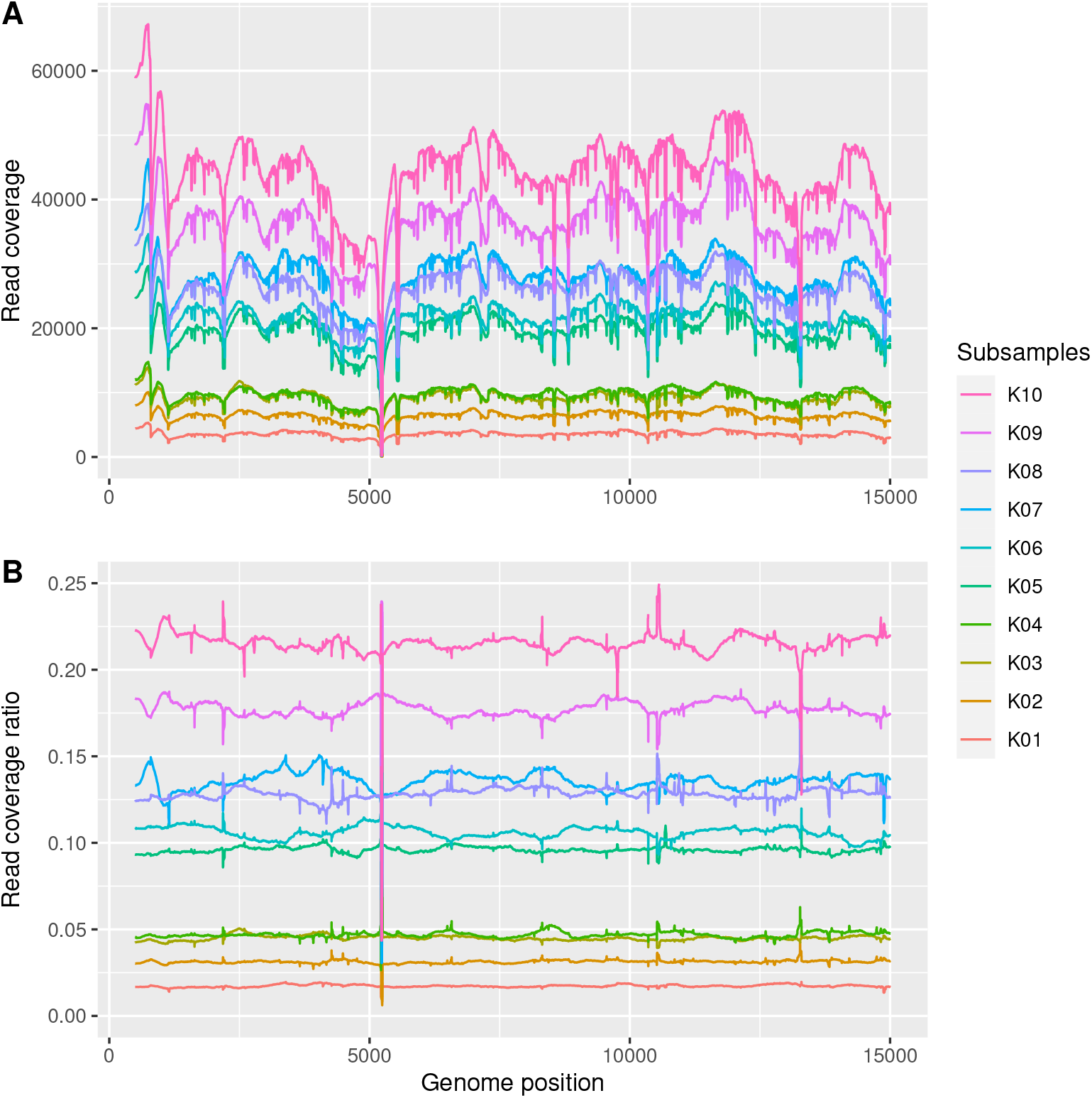
Distribution of the read alignments of 10 haplotypes against the reference genome. (A) The absolute read coverages along the genome. (B) The ratios on read coverages between sub-samples along the genome.

AFPhyloMix only considers the association between two SNP sites, but it is sensible to consider the association between more SNP sites in order to acquire higher sensitivity of the methods especially when the number of haplotypes increases. The time complexity of the algorithm is *O*(*mn^d^*) where *n* is the number of haplotypes, *m* is the number of potential SNP sites, and *d* is the number of SNP sites associated to construct patterns of nucleotides. In our current implementation of AFPhyloMix, *d* = 2. The running time of the algorithm will increase exponentially as *d* increases. It will be a challenge to come up with a faster algorithm and consider the association between more number of SNP sites, so that the method can work more effectively even for a mixture with a large number of sub-samples.

Finally, it is worth noting that we have applied AFPhyloMix to sequences of closely related individuals — in our simulations, we set a root-to-tip distance of 0.03. The assumption of an infinite sites model that is applied in AFPhyloMix is appropriate for closely-related individuals. Amongst other things, the infinite sites model allows us to constrain the number of site patterns we expect to see, and use deviations from these expected patterns to error-correct. To extend our algorithm to more divergent sequences will require a different model of mutation. This remains a work in progress.

## Conclusion

AFPhyloMix is designed to estimate the concentration of haplotypes and reconstruct the phylogenetic tree directly from the short read sequences in the mixture of haplotypes with no barcode, given that the number of haplotypes is known. This research demonstrates the feasibility of our approach, and is a first attempt to infer the phylogenetic tree from the mixture of reads unidentifiable to haplotypes, bypassing the assembly process of multiple genomic sequences. The experimental results have demonstrated that AFPhyloMix works reasonably well for both simulated data and real data.

## Supporting information

### S1 Appendix. Moves in Markov chain Monte Carlo

AFPhyloMix applies the Metropolis algorithm and proposes changes in either tip frequencies, error rate, or tree topology, serially. The following moves during the Markov chain Monte Carlo process are implemented. All these moves have Hastings ratio equal to 1.

- Update in tip frequencies For every tip *i* on the tree, its frequency (*f_i_*) is updated to *f_i_* + *r* where *r* ~ *U*(–0.15, 0.15). If the new value of frequency is a negative value, then the frequency is – (*f_i_* + *r*).
- Update in the error rate The error rate (*e*) is updated to *e* + *r* where *r* ~ *U*(–0.0005,0.0005). If the new value of error rate is a negative value, then the error rate is – (*e* + *r*).
- Distribute the frequencies between two tips Randomly select two tips *i* and *j* on the tree. Update their frequencies *f_i_* and *f_j_* to *r* and *f_i_* + *f_j_* – *r* where *r* ~ *U*(0, *f_i_* + *f_j_*).
- Swap the frequencies between two tips Randomly select two tips on the tree and swap their frequencies.
- NNI Randomly select an internal edge on the tree and perform Nearest-neighbor interchange (NNI).
- Swap between two subtrees Randomly select two nodes (internal or terminal) *A* and *B* satisfying the following criteria: (1) *A* and *B* are not sister nodes; (2) *A* is not an ancestor of *B*; and (3) *B* is not an ancestor of *A*. Then swap between the subtree rooted at *A* and the subtree rooted at *B*.
- Merge and split Randomly select an internal node *A* with exactly two leaves *i* and *j*. Select another tip *k* which is not a sister node of *A*. If *f_k_* > *f_i_*, then use the following step to merge the leaves *i* and *j* and split the leaf *k* into two tips. First remove the tip *i* and *j* and turn *A* into a new tip with frequency *f_i_* + *f_j_*. Then add two children at the node *k* with frequencies *f_i_* and *f_k_* – *f_i_*.
- Combine move Among the moves of NNI, Swap between two subtrees, and, Merge and split, randomly select two of them and consider to perform the two moves together.

#### Prior distributions

The prior of haplotype frequency is a gamma distribution with rate parameter 0.1 and shape parameter 2. The prior of the tree topology is uniform across all the possible topologies, and the prior of the error rate a uniform distribution with maximum value of *e_max_* (which is set to 0.01 for the reads produced from Illumina sequencing machines).

#### Metropolis-coupled Markov chain Monte Carlo

AFPhyloMix runs 8 Markov chain Monte Carlo processes in parallel: one cold chain and seven hot chains. The *i*-th hot chain’s temperature is set to (1 – *i*/8). A hot chain is randomly selected and its posterior probability is compared with that of the cold chain for every *x* iterations (where the value of *x* equals to the number of possible moves times 5, for example, *x* = 12 × 5 = 60 for 5 haplotypes; while *x* = 22 × 5 = 110 for 15 haplotypes). If the posterior probability of the hot chain is higher than that of the cold chain, two chains will be swapped. When swapping between two chains, we followed [29]’s implementation that the temperature of the two chains were exchanged instead of the states. Exchanging their temperatures are more efficient than exchanging their states, because the states which include many parameters are usually large in size.

## Availability of software and materials

The software AFPhyloMix and the materials are available in OSF repository: https://osf.io/w5s2h/ (DOI: 10.17605/OSF.IO/W5S2H)

## Acknowledgements

The authors thank David Bryant for constructive comments on the manuscript. This work was supported by computational resources provided by the Australian Government through the Australian National University under the National Computational Merit Allocation Scheme. This project was funded by an Australian Research Council Discovery Project Grant DP160103474.

